# Mapping brain volume changes in the zQ175DN mouse model of Huntington’s disease: a longitudinal MRI study

**DOI:** 10.1101/2025.11.04.686560

**Authors:** Nicholas Vidas-Guscic, Tamara Vasilkovska, Stefanie Pluym, Joëlle van Rijswijk, Johan Van Audekerke, Haiying Tang, Ignacio Munoz-Sanjuan, Dorian Pustina, Roger Cachope, Annemie Van der Linden, Daniele Bertoglio, Marleen Verhoye

## Abstract

Huntington’s disease (HD) is a progressive neurodegenerative disease affecting motor and cognitive abilities, as well as exhibiting psychiatric manifestations. Studies in people with HD (PwHD) consistently report atrophy of the caudate and putamen as an early pathological event and is therefore considered an early biomarker. Investigating whether similar phenotypic features are apparent in rodent HD models is important since it could have translational potential in evaluating the efficacy of novel therapeutic strategies. We used high-resolution anatomical images to longitudinally investigate brain morphology in the zQ175DN heterozygous mouse model (HET) and wildtype littermates (WT) at 3, 6, and 10 months of age (M), which reflect different stages of phenotypic progression. We investigated volumetric alterations using semi-automatic segmentations of HD on relevant regions-of-interest (striatum, cerebellum, corpus callosum, cerebral cortex, ventricles, and total brain volume) and whole brain voxel-wise Tensor Based Morphometry (TBM) analysis. The striatum showed the earliest progressive lower absolute volume in HET mice compared to WT, starting from 3M, followed by lower absolute volume of cortex and corpus callosum concomitantly at 6 and 10M. TBM highlighted lower relative local volume in the rostral-medial striatum at all ages, and in cerebral cortex in HET mice at 6 and 10M. A bigger relative local volume in the cerebellum was observed at all ages in HET mice, and in the globus pallidus, substantia nigra, at 6 and 10M. Overall, this study revealed key structural abnormalities that resemble the natural history of PwHD. Hence, analysis of brain structure through MRI in the zQ175DN heterozygous mouse model holds potential for testing disease-modifying treatments expected to slow down or prevent structural degeneration.

**Highlights:**

- The striatum of zQ175DN heterozygous mice shows volume decrease at 3 months compared to WT mice
- Widespread volume reductions are observed from 6 months in zQ175DN heterozygous mice
- Tensor Based Morphometry highlights vulnerability of the dorsomedial striatum in the zQ175DN heterozygous mouse model

## Introduction

Huntington’s disease (HD) is a progressive genetic neurological disease that comprises psychiatric manifestations as well as motor and cognitive impairments (Bates et al., 2015; Snowden, 2017). HD is caused by an expansion in the cytosine-adenine-guanine (CAG) trinucleotide repeat of the huntingtin gene (*HTT*), leading to the expression of toxic mutant huntingtin (mHTT) protein (MacDonald et al., 1993). The medium-sized spiny neurons of the striatum are most vulnerable to the cascade of neuropathological events in HD, and their loss is reflected in the macroscopic atrophy of the striatum (Fazl & Fleisher, 2018; Liu et al., 2023; Vonsattel et al., 1985; Wild et al., 2010) In addition, several other brain areas, such as cortex (Nopoulos et al., 2007; Rosas et al., 2005; Shishegar et al., 2019) and white matter (Paulsen et al., 2006; Tabrizi et al., 2009; Tabrizi et al., 2011) have been shown to be affected early on in people with HD (PwHD).

The clinical diagnosis of HD is based on established motor, cognitive, or behavioral signs (Tabrizi et al., 2022). However, pathological processes start many years before the onset of clinical signs and symptoms, and for this reason the Huntington’s Disease Integrated Staging System (HD-ISS) was recently proposed to include key events of HD progression. In HD-ISS, striatal atrophy serves as marker of Stage 1 progression given the strong effect and ubiquitous find in many HD studies (Tabrizi et al., 2022). Among the striatal structures, caudate is the first structure affected, followed by putamen and thalamus, a pattern that suggests a dorsoventral temporal gradient of atrophy (Ramirez-Garcia et al., 2020; Wijeratne et al., 2018). These findings are supported by local brain morphometry MRI studies using voxel-based morphometry (VBM) and tensor-based morphometry (TBM), which have revealed grey matter anomalies in the basal ganglia (Douaud et al., 2006; Kipps et al., 2005), which could already be detected in early stages (HD-ISS stage 1) of disease progression in PwHD (Tabrizi et al., 2011).

While efforts are ongoing to develop and test novel therapeutic strategies that delay or halt the progression of HD, a disease-modifying treatment has not been identified (Estevez-Fraga et al., 2020; Ferguson et al., 2022). Biomarkers are needed to identify specific anomalies or their mitigation post-intervention (Zeun et al., 2019). To evaluate the therapeutic efficacy of new and promising drugs using established biomarkers, such as striatal atrophy —a key biomarker in HD— identifying comparable biomarkers in animal models may play a crucial role as phenotypic development occurs at a shorter time scale (Mukherjee et al., 2022).

The zQ175DN HET mouse model is a genetic model of HD with a human HTT exon 1 with ∼190 CAG repeats knocked into the murine *Htt* gene. Since zQ175 is already well studied for its phenotypic and neuropathologic similarity to human disease, this model is proposed as an ideal candidate to investigate potential biomarkers and therapies. This mouse model exhibits striatal transcriptional and proteomic expression dysregulation, similar to PwHD (Atherton et al., 2016; Callahan et al., 2022; Deng et al., 2021; Goodliffe et al., 2019; Herrmann et al., 2021; Langfelder et al., 2016; Lee et al., 2020; Smith et al., 2014). zQ175 mice exhibit progressive striatal abnormalities, i.e., HTT aggregate pathology starting in the caudate putamen at 3 months (M) and later in the cortex at 8M (Carty et al., 2015).

Characterization of the zQ175DN HET mice along phenotypic progression has already revealed changes in brain perfusion and dynamic functional brain alterations as early as 3M, and develops a more complex pathology as the phenotype advances (Adhikari et al., 2023; Vasilkovska et al., 2023; Vasilkovska, Salajeghe, et al., 2024; Vasilkovska, Verschuuren, et al., 2024). Importantly, white matter characterization in this model shows microstructural anomalies in the striatum, which are already present at 3M, extending to other white matter structures such as the corpus callosum as mice become older (Vidas-Guscic et al., 2024). However, a detailed assessment of the volumetric changes along phenotypic progression in the zQ175DN HET mouse model is still lacking.

Consistent reductions in brain volume provide a strong foundation for applying volumetric MRI and voxelwise TBM as translational tools for biomarker discovery. Compared to conventional region-of-interest (ROI) delineation or template-based brain volume assessment (Heikkinen et al., 2012; Peng et al., 2016), TBM allows for more sensitive, whole-brain voxel-wise assessment of regional atrophy patterns over time, allowing the detection of subtle and progressive structural changes with age. By integrating TBM with high-resolution (78 µm, isotropic) three-dimensional (3D) MRI in the zQ175DN model, it becomes possible to map the spatio-temporal trajectory of brain structural changes, identify early imaging markers, and quantify treatment effects in preclinical interventional studies. The zQ175DN HET mouse line represents a potential platform to evaluate not only functional and molecular but also structural neuroimaging-based biomarkers in HD research. Our study employs an extensive ROI analysis complemented by a voxel-based TBM analysis, which assesses local deformation relative to a healthy control template. Hereby, TBM can be used to pursue an unbiased analysis of any region that could possess biomarker characteristics independently of prior hypotheses.

The objective of this study was to investigate volumetric anomalies in the zQ175DN HET mouse model at 3, 6 and 10M, compared to WT littermates. We followed this objective through longitudinal measurements from the same animals and adopted TBM in conjunction with ROI analyses to detect potential biomarkers of disease progression in HET mice. Based on prior knowledge of human and animal studies, we hypothesized that the zQ175DN HET mouse model will exhibit progressive volumetric anomalies in HD-relevant regions, specifically in the striatum and cortex.

## Material and methods

### Animals

Age-matched male zQ175DN heterozygous (HET) knock-in mice (N=18) and zQ175DN wildtype littermates (N=18) (JAX stock #029928) were obtained from the Jackson Laboratory (Bar Harbor, ME, USA). The zQ175DN HET mouse model is a subtle modification of the original zQ175 mouse model in which the 5’ neomycin cassette was removed (Farshim & Bates, 2018; Heikkinen et al., 2012; Menalled et al., 2012) and has the human HTT exon 1 substitute for the mouse Htt exon 1 with ∼180–220 CAG repeats long tract. zQ175 mice show circadian dysfunction, amotivation, cognitive impairment, and gait abnormalities (Beaumont et al., 2016; Covey et al., 2016; Heikkinen et al., 2020; Oakeshott et al., 2012; Piiponniemi et al., 2018; Smarr et al., 2019; Smith et al., 2014).

Due to sporadic congenital portosystemic shunt occurring in C57BL/6J mice (Cudalbu et al., 2013), all animals were screened at Jackson Laboratories before shipment to be shunt-free. Mice were genotyped at birth and after experiments to determine their number of CAG repeats. Body weights of the mice were measured at the ages of 3M, 6M, and 10M (**Suppl. Fig. 1**). Based on our prior experience with this model, all animals were single-housed at the central housing facility of the University of Antwerp to prevent the potential development of aggressive dominant-submissive behavior. All animals were housed on a 12-hour light-dark cycle in climate-controlled rooms. Animals were provided with nesting material, food and water *ad libitum*. All experimental procedures and animal handling described in this investigation were carried out according to European guidelines for the housing and use of laboratory animals (2010/63/EEC). This project was approved by the Ethical Committee on Animal Care and Use at the University of Antwerp, Belgium (ECD 2017-09).

### MRI acquisition

MRI acquisition was performed for all mice at the age of 3M, 6M and 10M on a 9.4 Tesla horizontal preclinical BioSpec MRI scanner equipped with a volume RF transmit coil and a four-element receive-only mouse head CryoProbe RF receive coil (Bruker Biospin MRI, Ettlingen, Germany). The 3D anatomical images were acquired within a longer scan protocol for functional MRI in the resting state functional MRI (rs-fMRI) (see Vasilkovska et al., 2023), which required the following anesthesia regimen: mice were initially anesthetized with 2% isoflurane (Isoflo®, Abbot Laboratories Ltd., USA) in a mixture of 200 ml/min O_2_ and 400 ml/min N_2_. Next, mice were injected with a bolus of medetomidine (0.075 mg/kg; Domitor, Pfizer, Karlsruhe, Germany), while isoflurane levels were lowered to 0.4% for rs-fMRI acquisition. After 60 minutes, the medetomidine infusion was stopped, and isoflurane was increased to 1-2% directly before structural imaging. Throughout the duration of the experiment, all physiological parameters (breathing rate, heart rate, O_2_ saturation, and body temperature) were monitored. The experiments were continuously alternated between WT and HET mice, with a per group randomization of mice for each acquisition at each age.

Prior to image acquisition, a B_0_ field map was acquired to measure magnetic field inhomogeneities, which was followed by local shimming within an ellipsoid covering the brain.

For structural imaging, a T_2_-weighted 3D turboRARE sequence was used with following parameters: Field of view (20×20×10)mm^3^, image matrix [256×256×128], voxel size (78×78×78)µm^3^, repetition time 1800ms, echo time 42ms, RARE factor 16, partial-Fourier acceleration of 1.4 in slice-phase encoding direction requiring an effective acquisition matrix of [256×256×91]. The total scan duration of the T_2_w acquisition per animal was 43 minutes and 41 seconds.

### Image preprocessing

All data was blinded to the researchers to avoid subjective decision making during manual processing steps. T_2_-weighted 3D turboRARE images were first debiased using the N4biasfieldcorrection algorithm (ANTs, https://stnava.github.io/ANTs/). To facilitate skull stripping of all the 3D data, an intermediate template was generated from the T_2_-weighted debiased 3D turboRARE images from all 36 subjects at all ages. All T_2_-weighted debiased 3D turboRARE images were registered to this intermediate study-based template, using affine transformation, and non-linear warps (ANTs). A whole brain mask was created in study-based template and backtransformed using the inverse operation of the affine transformations and non-linear warps from the aforementioned subject to the template space registration, using nearest neighbor interpolation (ANTs). The resulting whole brain masks in subject space were visually checked and manually corrected if necessary and then used for skull stripping.

An updated final template was generated using the skull stripped images of 3M old WT subjects, which was used as target space for all further analysis and hereafter referred to as the WT template. Next, all individual skull stripped T_2_-weighted debiased 3D turboRARE images were normalized to the WT template using affine transformation and non-linear warps (ANTs). The resulting non-linear warps were used for further TBM analysis. For a visual representation of the processing and analysis workflow, see **supplementary figure 2**.

### Whole-brain and ROI volume analysis

To assess large-scale macroscopic brain structural alterations, voxel counts were extracted from the back transformed and manually corrected whole brain masks, the cerebrum and cerebellum. The striatum, cerebral cortex, corpus callosum, and ventricles were selected for a localized ROI-based volume analysis based on prior reports of volumetric alterations in these regions (Reiner et al., 2011; Rosas et al., 2010). These ROIs were first manually delineated on the WT template in Mrview (Mrtrix3, https://www.mrtrix.org/) following anatomical parcellations in the Allen Mouse Brain atlas (Allen Institute for Brain Science (2004), Allen Mouse Brain Atlas available at (https://mouse.brain-map.org/). A parcel of the cerebrum was created by subtracting the cerebellum from the whole brain parcel.

Next, parcels were then retransformed using the inverse operation of the registration of each subject in native space to the WT template. The quality of the transformed parcellations in the native space was visually assessed and adjusted if necessary. Next, voxel counts and absolute volume (in mm^3^) of the ROIs were extracted for each parcel.

### Tensor-Based Morphometry Analysis

Voxel-based-TBM analysis was used to identify regions that show genotypic differences in the relative local deformations to a reference target per age. The aforementioned symmetric non-linear (SyN, ANTs) warps from the WT template to each T_2_-weighted 3D turboRARE image were used to calculate the Jacobian determinants using ANTs software. The Jacobian determinant describes only local deformations since global size differences (i.e., if brains were of different sizes) were corrected by the preceding affine registration step. Forward non-linear warps describe the behavior of the WT template (fixed image) to the individual 3D T_2_-weighted turboRARE (moving image). Therefore, Jacobian determinant values <1 indicate a shrinkage of the template to fit with the subject image and can be interpreted as a smaller local area of the subject image. On the opposite end, Jacobian determinant values >1 can be interpreted as larger local areas in the subject images. Jacobian determinant values were log-transformed to improve statistical analysis (West, 2022).

An ROI-based analysis was performed to investigate genotype x age interactions of the average log-transformed Jacobian determinant guided by the observations of the voxel based TBM analysis. These ROIs included the striatum, globus pallidus, substantia nigra, cerebellum, cerebral cortex, and hippocampus. For each ROI parcel, the ROI-averaged values of the log-transformed Jacobian determinants were extracted using MRtrix3.

### Statistical analysis

To assess the effect of genotype across different time points, a linear mixed model (LMM) was performed on the absolute (total) volume of the whole brain and ROIs expressed in mm^3^, and ROI based log(Jacobian determinant), where a main effect of zQ175DN genotype (WT and HET) and age (3M, 6M, 10M) and the (age x genotype) interaction was tested. In case of an interaction effect, a post hoc t-test was performed to test for age-specific genotype effects and group-specific age affects (p<0.05, False Discovery Rate (FDR) corrected). To test for differences in the voxel-based TBM metrics between the two zQ175DN genotype at the different ages (3M, 6M, and 10M), unpaired t-tests were performed using permutation testing and Threshold Free Cluster Enhancement (TFCE) followed by Family Wise Error (FWE) correction (Winkler et al., 2014).

## Results

### Volumetric decreases originate in the cerebrum in zQ175DN HET mice

A linear mixed model was used to assess the changes in global brain volume in relation to genotype and aging. The LMM results showed a significant interaction effect (Suppl. table 1). Post hoc testing (Suppl. table 1) revealed a statistically significant difference in total brain volume between the genotypes at 6M and 10M, which emerged due to an excessive decrease in total brain volume in zQ175DN HET compared to WT between 3M and 6M of age (Fig. 1A). While the whole brain volume remained lower after 6M in zQ175DN HET, the WT mice exhibited an increase in whole brain between 6M and 10M. The whole brain differences were found to be driven by the cerebrum, as the cerebrum volume demonstrated similar differences between HET and WT mice (**Fig. 1B**), while the volume of the cerebellum showed only a main aging effect without any interaction with the genotype (**Fig. 1C**).

**Figure 1:**
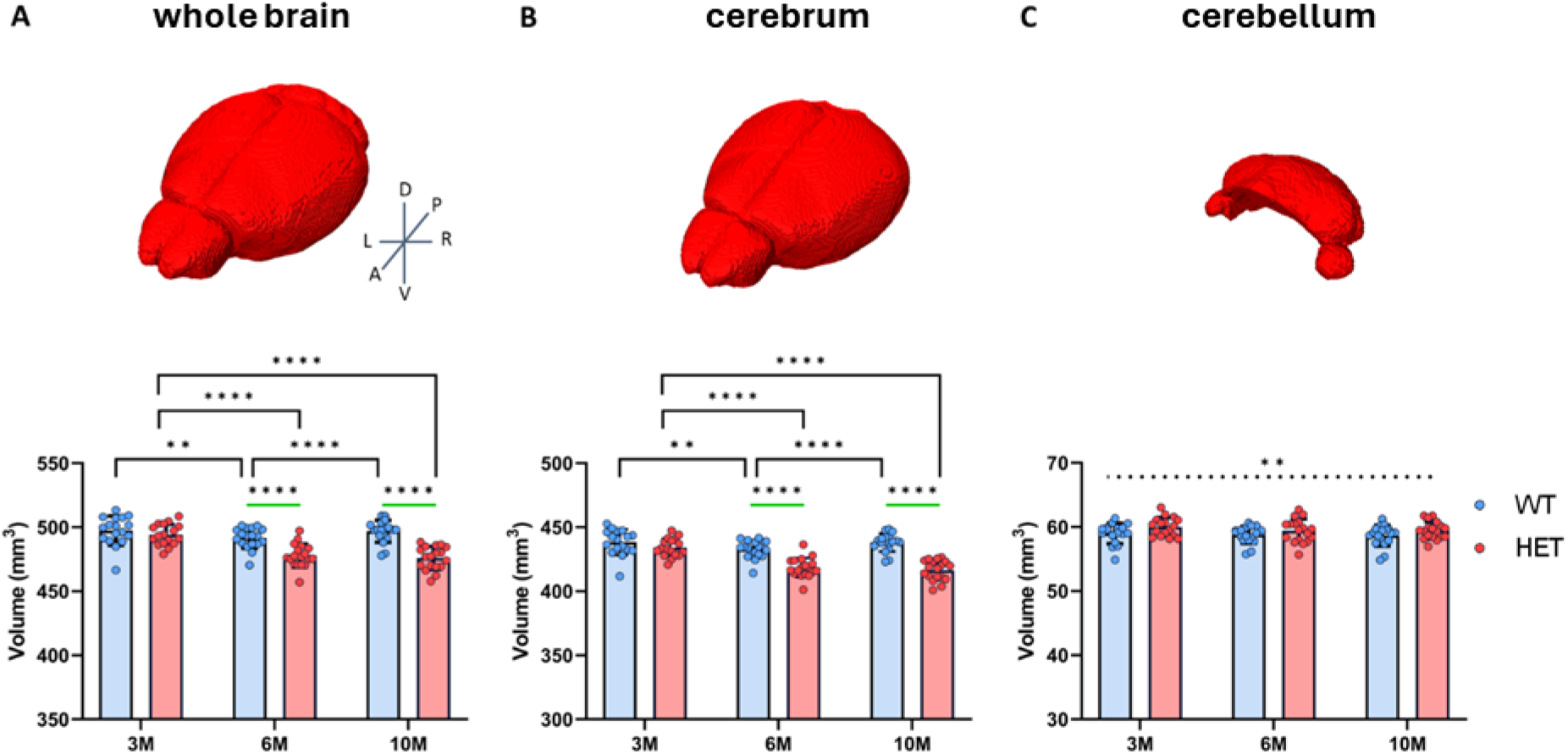
Absolute volume of whole brain, cerebrum, and cerebellum of zQ175DN HET and WT animals at 3, 6, and 10M. Three-dimensional volume rendering and graph of absolute volumes of WT (blue) and HET (red) for (**A**) whole brain volume, (**B**) cerebrum, and (**C**) cerebellum. The outcome of the LMM with FDR correction and post hoc tests is indicated as follows: green lines indicate post hoc genotype effects, black brackets indicate post hoc age effects and horizontal dashed line indicates a main age effect. ** p < 0.01, **** p < 0.0001. Error bars represent *SD*.

### Basal ganglia related structures show volume anomalies in zQ175DN HET mice

Next, the volumes of different HD-relevant regions were investigated using a linear mixed model (**Fig. 2; Suppl. table 2**). For the striatal volume, a statistically significant interaction was observed. The results of the post hoc testing are summarized in **Suppl. Table 2**. These tests revealed smaller striatal volume in HET compared to WT mice at 3M, 6M, and 10M. In HET mice, a gradual decrease in striatal volume was observed between all investigated ages, while in WT mice only a slight decrease in striatal volume happened from 3M to 6M.

**Figure 2:**
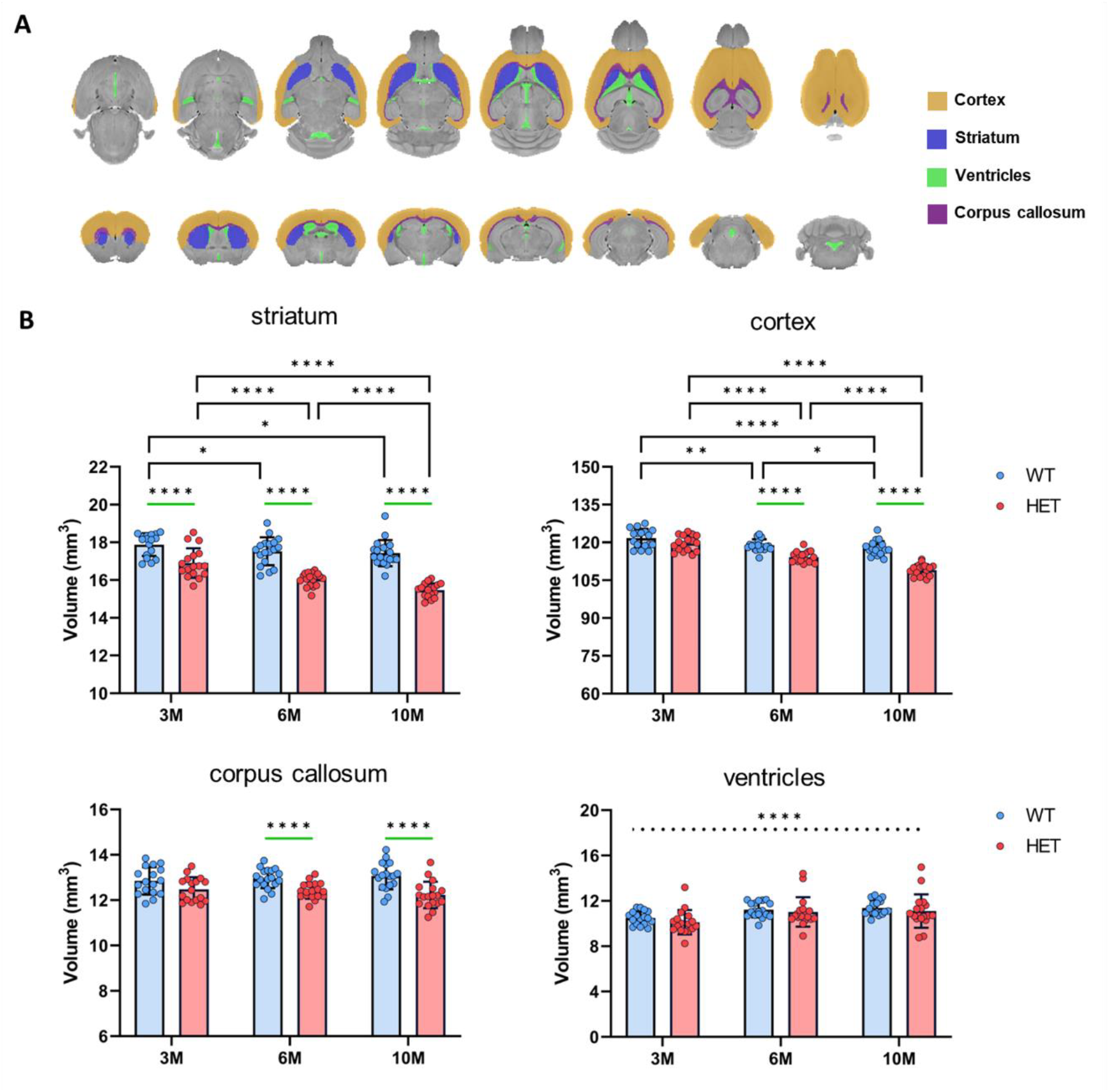
Absolute volume quantification of ROIs of zQ175DN HET and WT animals at 3, 6, and 10M. (**A**) Parcellations of ROIs for volume extraction on the T_2_-weighted 3D-turboRARE template. (**B**) Graphs of absolute volumes of WT (blue) and HET (red) for striatum, cortex, corpus callosum and ventricles. The outcome of the LMM with FDR correction and post hoc tests is indicated as follows: green lines indicate post hoc genotype effects, black brackets indicate post hoc age effects. Horizontal dashed line indicates a main age effect. * p<0.05, ** p < 0.01, *** p < 0.001, **** p < 0.0001. Error bars represent *SD*.

In the cortex, a significant interaction was observed. A significantly lower volume of the cortex was observed in HET compared to WT mice at 6M and 10M. This was driven by an age-wise decrease that was greater in HET mice compared to WT mice at all ages.

In the corpus callosum, a significant interaction was observed. Post hoc tests revealed significantly lower absolute volume of the corpus callosum in HET mice compared to WT mice at 6M and 10M (**Suppl. table 2**).

The volume of the cerebral ventricular system did not show significant interaction, but a main effect of age was observed as the ventricles become larger with age (Fig. 2B; Suppl. table 2).

### TBM reveals relative local deformations in disease-relevant brain structures in zQ175DN HET mice

In this study, cross-sectional comparisons using two-sample t-tests were performed to assess age-related genotypic differences in local deformations (**Fig. 3**). At 3M, a small cluster in the cerebellum showed a larger relative local volume in HET mice compared to WT mice (**Fig. 3**). At 6M, a significantly lower relative local volume was observed in the dorsomedial striatum, along with a significantly higher relative local volume in the white matter/arbor vitae of the cerebellum in HET mice compared to WT mice. At 10M, the differences were more extensive, with a significantly lower relative local volume in the striatum, cortex, and hippocampus in HET compared to WT mice, and higher volume in the arbor vitae of the cerebellum, the globus pallidus and substantia nigra (**Fig. 3**).

**Figure 3:**
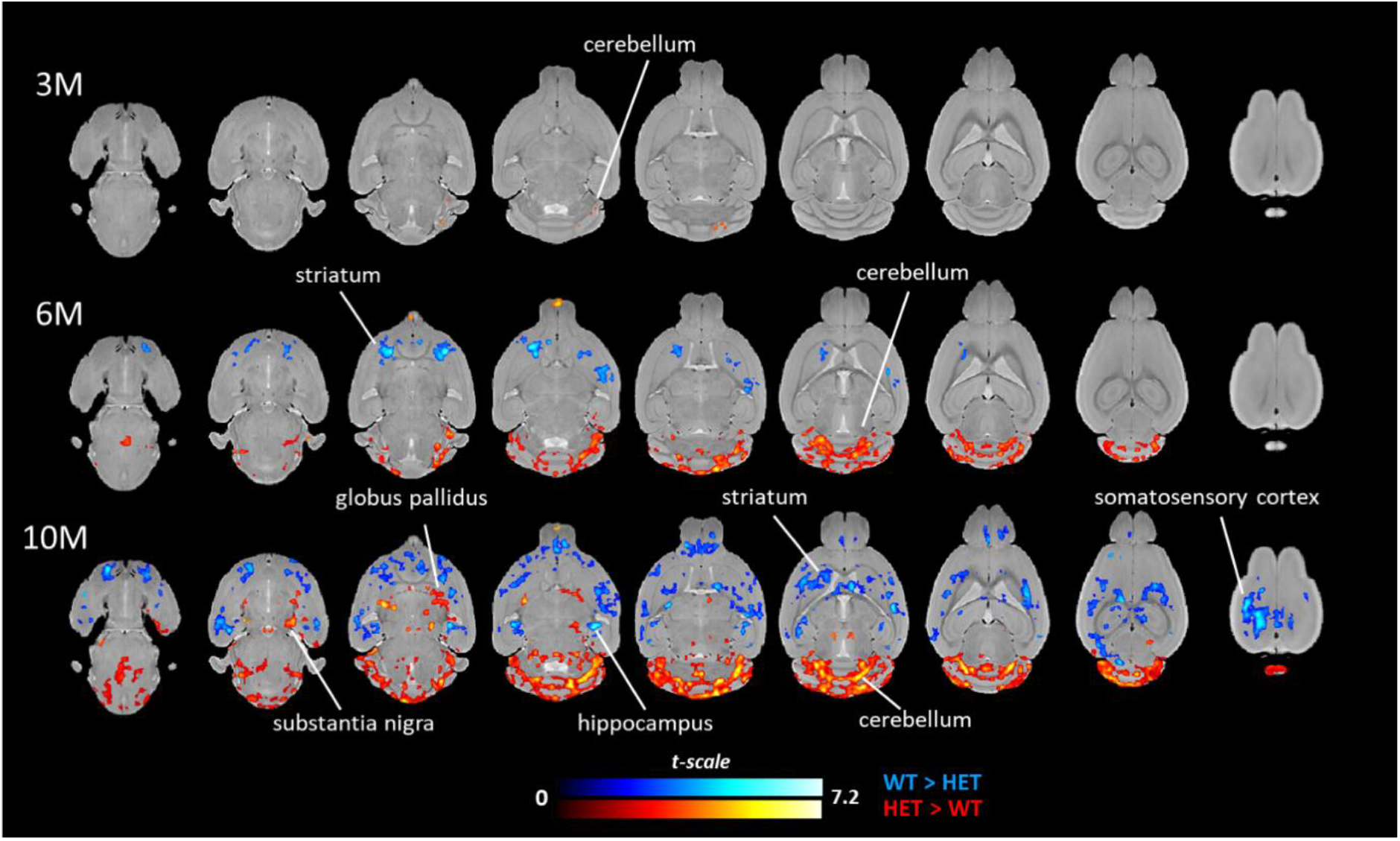
Difference in relative local deformations between zQ175DN HET and WT animals using TBM at 3, 6 and 10M. Statistical t-map (P_TFCE-FWE_ < 0.05) of two-sample t-tests showing regions presenting a significant genotype effect at 3M (top row), 6M (middle row) and 10M (bottom row), overlaid on the WT template. Colors indicate results of *t*-statistics with blue/red colors indicating voxels that have a significant lower relative log(Jacobian determinant)/higher relative log(Jacoban determinant) in HET compared to WT mice, respectively.

To investigate whether the increased relative local volume in the cerebellum of HET mice was not solely caused by the significantly higher whole brain affine scaling factor in HET compared to WT mice, separate additional TBM analyses were performed on 1) cerebrum and 2) cerebellum separately. First, the affine scaling factor was calculated from the single-vector decomposition (SVD) of the affine transformation matrix. The affine scaling factors extracted for both the whole brain analysis and cerebrum masked analysis revealed a significant interaction with post hoc test indicating a significantly lower scaling factor for HET compared to WT mice at 3M, 6M and 10M (Suppl. Fig. 3). In contrast, no significant differences were observed in the scaling factor of the cerebellum masked analysis (**Suppl. Fig. 3**). This is in line with absolute volume analysis of the whole brain, cerebrum and cerebellum as shown in **Fig. 1**. The additional TBM analyses showed that the same regions and direction of the effects for the relative local deformations are present in the cerebrum (**Suppl. Fig. 4A**) and cerebellum (**Suppl. Fig. 4B**) masked TBM analysis as the whole brain TBM analysis with a smaller number of voxels showing significant genotype differences.

Next, we performed a statistical analysis of the ROI-based averaged relative local deformations for which HD-relevant ROIs were selected (striatum and cortex) and additional ROIs that showed genotypic differences in the whole brain TBM analyses per age (globus pallidus, substantia nigra, hippocampus, and cerebellum) **Fig. 4**.

**Figure 4:**
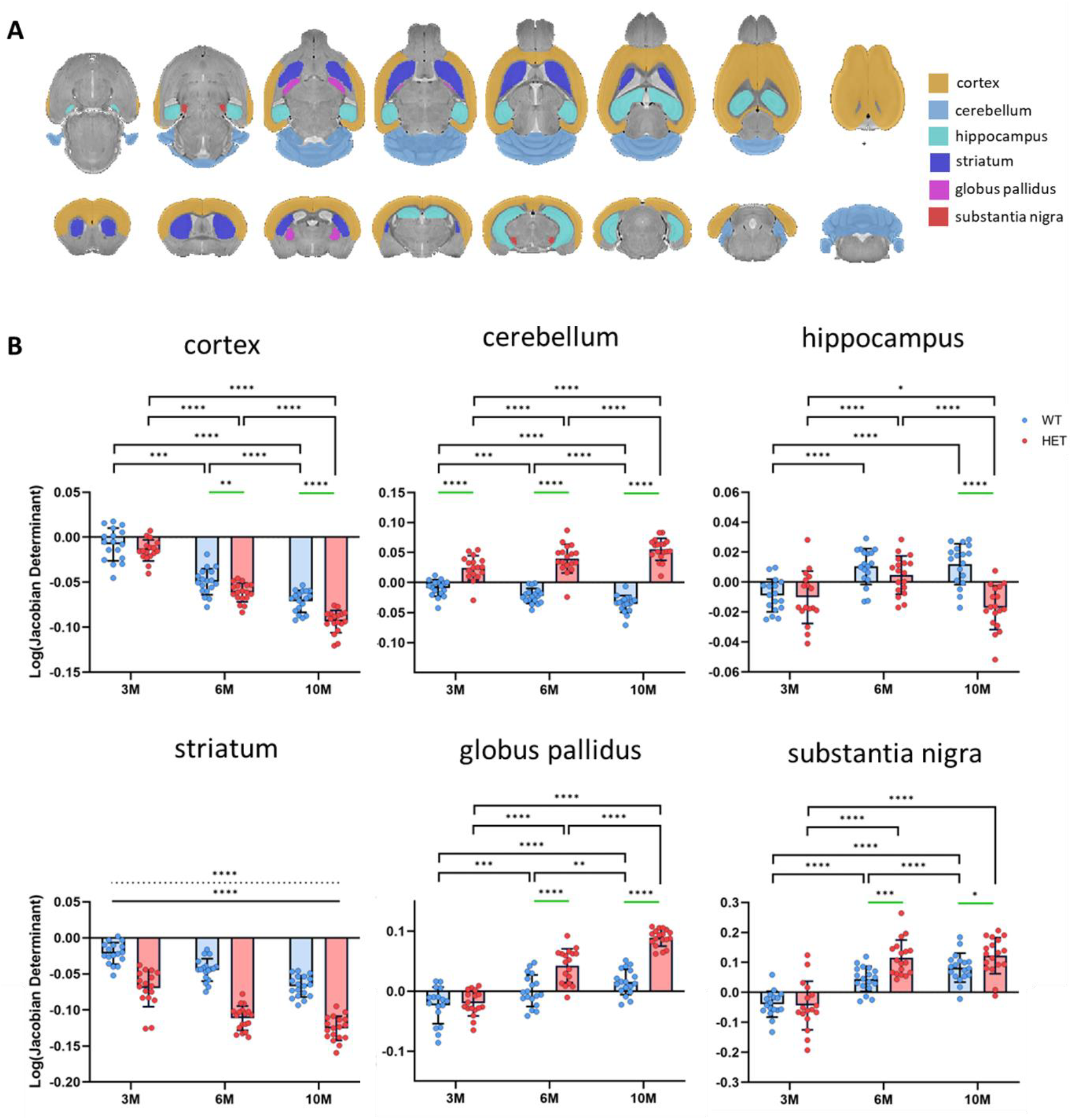
Comparison of average local deformations in ROIs of zQ175DN HET and WT animals. (**A**) Parcellations of ROIs guided by TBM analysis for average log(Jacobian determinant) extraction overlaid on a T_2_-weighted 3D-turboRARE in two orthogonal views. (**B**) Graphs of average log(Jacobian determinant) of WT (blue) and HET (red) for the cortex, cerebellum, hippocampus, striatum, globus pallidus and the substantia nigra. The outcome of the LMM with FDR correction and post hoc tests is indicated as follows: green lines indicate post hoc genotype effects, black brackets indicate post hoc age effects. Horizontal solid line and dashed line indicate main genotype and age effects, respectively. * p < 0.05, ** p < 0.01, *** p < 0.001, **** p < 0.0001. Error bars represent *SD*.

Significant interaction effects were observed for all regions, except for the striatum which showed both a significant genotype and age effect (**Suppl. table 3**). For the cortex, post hoc tests indicated a significantly lower relative local volume in HET mice compared to WT mice at 6M and 10M, driven by a more pronounced decrease from 3M to 10M in HET mice compared to WT mice (**Suppl. table 2**).

In the cerebellum, post hoc tests showed that HET mice have significantly higher relative local volumes compared to WT mice, driven by a progressive volume increase in HET from 3M to 10M, while WT mice showed a relative volume decrease (**Suppl. table 4**).

For both the globus pallidus and the substantia nigra, post hoc tests revealed a significantly higher relative local volume in HET compared to WT mice at 6M and 10M. A progressive increase in the globus pallidus was observed in both genotypes, while in the substantia nigra, HET mice had significant volume increases from 3M to 6M and 3M to 10M, unlike WT mice, which increased continually from 3M to 10M (**Suppl. table 4**).

In the hippocampus, post hoc testing revealed a significantly smaller relative local volume in HET compared to WT mice at 10M. The relative local volume in the hippocampus of HET mice did not follow the ageing effect that was observed in WT mice. HET mice showed a decrease from 6M to 10M, while no significant change was observed in WT mice from 6M to 10M. (Suppl. Table 4).

## Discussion

This study aimed to characterize the volumetric changes that occur during phenotypic progression in the zQ175DN HET mouse model of HD. We hypothesized that the zQ175DN HET mouse model demonstrates progressive volume loss, first in the striatum before affecting other regions of the brain. We demonstrated that volumetric anomalies can be detected in the striatum at the earliest time point investigated (3M) while the whole brain volume is still similar between genotypes. However, from 6M of age, the differences between the genotype groups were more widespread involving the whole brain, the cortex and the corpus callosum, while the cerebellum and the ventricular system remained relatively unaffected. Using TBM, a method that allows for data-driven (parcellation-free) characterization of local volume changes, we were able to detect localized subregional alterations in multiple brain regions.

This study reports the first evidence of striatal volume reductions in zQ175DN HET mice, observable at 3M before the bulk of local alterations which occur in other brain regions starting at 6M, resulting in detectable whole brain volume reductions. A division of the brain into the cerebrum and cerebellum suggests that whole brain anomalies are driven by alterations in the cerebrum, whereas the cerebellum volume remains unaffected by disease processes. Other studies investigating the Q175 HET model have not observed whole brain volume anomalies at 3M (Bertoglio et al., 2018; Heikkinen et al., 2012). A possible explanation for this discrepancy is the resolution of the structural MRI data used. Our study acquired three-dimensional data at a much higher resolution (78×78×78) µm^3^ which allows for a more precise delineation of the brain, thereby improving the sensitivity of our readouts.

Nevertheless, previous studies in the Q175 HET model have consistently reported abnormally low striatal volume (Bertoglio et al., 2018; Heikkinen et al., 2012, Heikkinen et al., 2021; Tereshchenko et al., 2019), with the earliest reported volumetric deficits in the adult stage of Q175 HET mice at 4M (Heikkinen et al., 2012). Our results suggest that, in this model, striatal atrophy can be detected even earlier, at 3M, which is the earliest timepoint in which a lower striatal volume has been detected in adult mice.

Prominent striatal vulnerability, expressed as a volumetric reduction compared to age-matched wildtypes, is in line with other MRI studies of mouse models of HD. Characterization of the more severe zQ175 homozygous mouse model showed that striatum, cortex, and whole brain volumes are significantly reduced 3, 6, 9 and 12M (Peng et al., 2016), while the slower disease phenotype progression of the Q140 model leads to volume reductions at a later age (18M) (Pérot et al., 2022).

As the striatum is a functionally segregated region, it is important to understand which compartments of the striatum are earliest affected (Foster et al., 2021; Hintiryan et al., 2016). Using TBM, starting at 6M of age, we observed decreased local volume in the dorsal striatum and, more prominently, in the dorsomedial portion. In line with these findings, our previous work utilizing diffusion MRI documented changes in white matter properties starting at 6M of age in the zQ175DN HET mouse model, where a reduction in fiber cross-section was also observed in the dorsomedial striatum (Vidas-Guscic et al., 2024).

The most abundant population of neurons in the striatum, the medium-sized spiny neurons, are selectively vulnerable in HD patients (Ehrlich, 2012), leading to hallmark degeneration of the caudate and putamen (Tabrizi et al., 2022). However, there is no evidence that the observed volumetric deficits in the zQ175DN HET mouse model are the result of cell death, as no cell death has been reported in the striatum up to 15 months of age, despite the high CAG repeat length in this model (Deng et al., 2021; Ibrahim et al., 2023). Other factors must contribute to the observed striatal volume loss present at these early ages. For example, it is possible that the size or shape of cells is altered, i.e., in the Q175 mice D1 medium-sized spiny neurons show changes in dendritic arbors and reduced spine density (Goodliffe et al., 2019; Indersmitten et al., 2015). Other cell types besides neurons can contribute to these volumetric changes, too. A longitudinal characterization of astrocytes and pericytes, including changes in the deposition of mHTT in these cells, has shown that, starting at 6M, there is a marked increase in mHTT deposition in both astrocytes and pericytes, specifically in the dorsomedial portion of the striatum, aligning with the MRI structural changes observed (Vasilkovska, Verschuuren, et al., 2024). Despite no overt cell loss, these volumetric changes can represent early neurodegenerative remodeling, which constitutes retraction of neuronal dendrites and synapses, reduced astrocytic protrusions and microvascular pruning processes. Altogether, these factors can produce a compaction of the striatal architecture that is sufficient to be detected with MRI-based volumetric measures.

In our study, volume decreases in the corpus callosum and external capsule system were observed in the zQ175DN mice, starting at 6M. This finding aligns with our previous longitudinal study that characterized microstructural properties in the corpus callosum in the same animal model. Specifically, we previously observed decreases in both fiber density and cross-section at 6M, indicative of axonal loss and sparsity of the remaining white matter (Raffelt et al., 2017; Vidas-Guscic et al., 2024). Notably, white matter volume loss and white matter tissue microstructural anomalies have been reported in other mouse models of HD as well as human studies (Casella et al., 2020; Fraga et al., 2020; Gregory et al., 2018; Klöppel et al., 2008; Rosas et al., 2018; Tabrizi, Scahill, Durr, Roos, Leavitt, Jones, Landwehrmeyer, et al., 2011).

A lower cortical volume was observed at 6M, which has been reported to be significantly lower in Q175neo mice at 4M (Heikkinen et al., 2012). Cortical thinning is a feature of HD and mainly occurs in the motor cortices, followed by the parietal cortices (Ramirez-Garcia et al., 2020). In contrast, the absolute volume of the cerebellum was not significantly different between HET and WT mice in our study. This is consistent with reports in the Q175 mouse model of HD, other mouse models of HD (Carroll et al., 2011; Tereshchenko et al., 2019) and human structural MRI studies (Paulsen et al., 2010; Tereshchenko et al., 2019). While the total volume of the cerebellum is not significantly altered in the zQ175DN HET mouse model, we observed relative local volumetric increases assessed using TBM.

Such local volume increases could be the result of global affine scaling prior to non-linear image registration to fit the WT template. In this study, the affine scaling factor is mainly driven by the cerebrum which comprises 90% of the total brain volume, whereas the global scaling of the cerebellum is not significantly different between the genotypes. Such effects could potentially lead to compensatory non-linear warping to match the subject image with the template image causing relative local volumetric increases in cerebellum. Though the affine scaling factor of the cerebellum-masked TBM analysis did not present any significant genotype differences at any age, we could still observe significant increases in local volume of the cerebellum, collectively suggesting that significant local volumetric increases in the cerebellum of HET mice are due to local deformations rather than solely a consequence of affine scaling. Accumulation of mHTT aggregates occurs mainly in the granular and Purkinje cell layer (Rüb et al., 2013), which comprises a small portion of the cerebellum. Hence, alterations in this layer might not be detected in the structure as a whole. This could explain why a more sensitive measure of local deformations like TBM picks up alterations which are not observed in the total cerebellar volume.

Using TBM analysis, we also found local deformations within the known HD-relevant structures including the basal ganglia (striatum, globus pallidus, substantia nigra). The TBM analysis revealed larger relative local volume in the globus pallidus and substantia nigra of zQ175DN HET mice compared to WT littermates. Q175 mice have more abundant enkephalin immunostained terminals in globus pallidus externus from 6 months onward. The substance P immunostained terminals from striatal direct pathway neurons are also more abundant in the globus pallidus internus and substantia nigra from 6 months onward (Deng et al., 2021). This increased spiny projection neuron innervation could reflect a compensatory mechanism, for the reduced inhibition by the spiny projection neurons and explain the local relative volume increase in HET compared to WT mice observed in the globus pallidus and substantia nigra observed using TBM.

Ventricular enlargement is commonly reported in PwHD (Hobbs et al., 2010), but was not observed in this study. The gap in findings may certainly derive from different neuropathologies between species (i.e., loss of neurons in humans and no loss of neurons in mice) but also because we investigated only male zQ175DN HET mice. Lastly, the exact molecular mechanisms driving neuronal atrophy are not yet fully understood, therefore it is not known what is driving the volumetric alterations in this mouse model. In contrast to HD in humans, cell loss is not the driver of the volumetric changes in this model at the ages that are investigated. Further studies are required to investigate the tissue characteristics of the zQ175DN HET mouse model in more detail.

The results of this study have enhanced our understanding of the progression of volumetric alterations in the zQ175DN HET mouse model. The early longitudinal striatal volumetric reduction preceding more widespread structural alterations resembles the phenotypic progression in PwHD and could therefore hold potential as an early marker for disease progression and to investigate disease-modifying treatments using the zQ175DN HET mouse model.

## Conclusion

We demonstrated that the zQ175DN HET model undergoes morphological changes through age that are in line with those observed in human studies and follow a longitudinal worsening. Similar to the human situation, the striatum seems to be most vulnerable to the toxic effects of mHTT, showing early volumetric reduction preceding more widespread structural alterations. TBM confirms these findings and highlights the dorsomedial striatum as subregion of the striatum with increased vulnerability. Therefore, TBM could provide a candidate marker to assess the efficacy of HD-modifying therapeutics and to monitor disease progression.

## Data availability

Data presented in this paper is available in BIDs format upon reasonable request by contacting the corresponding author.

