## Supplementary table 1 for "Mapping brain volume changes in the zQ175DN mouse model of Huntington’s disease: a longitudinal MRI study"

**Supplementary table 1:** Outcome of linear mixed model to test the main effects of age and genotype and (age x genotype) interaction on the absolute volumes of the whole brain, cerebrum and cerebellum ($p < 0.05$). Post hoc testing (FDR corrected, $p < 0.05$) was performed in case of significant (age x genotype) interaction. Red font indicates a significant effect.

| | | Whole brain | | | Cerebrum | | | Cerebellum | | |
|---|---|---|---|---|---|---|---|---|---|---|
| **LMM effects** | | **DF** | **F-value** | **P-value** | **DF** | **F-value** | **P-value** | **DF** | **F-value** | **P-value** |
| | **Genotype** | 1.67, 55.19 | 103.30 | 0.0001 | 1.72,56.72 | 106.70 | 0.0000 | 1.94,60.55 | 5.30 | 0.0825 |
| | **Age** | 1,34 | 19.28 | 0.0000 | 1,34 | 30.39 | 0.0048 | 1,34 | 3.20 | 0.0091 |
| | **Genotype*age** | 2,66 | 57.91 | <0.0001 | 2,66 | 63.05 | <0.0001 | 2,66 | 1.06 | 0.3513 |

| | | Whole brain | | | | Cerebrum | | | | Cerebellum | | | |
|---|---|---|---|---|---|---|---|---|---|---|---|---|---|
| **post-hoc effects** | | **mean dif** | **t-value** | **P-value** | **DF** | **mean dif** | **t-value** | **P-value** | **DF** | **mean dif** | **t-value** | **P-value** | **DF** |
| **WT -HET** | **3M** | 3.28 | 0.97 | 0.3852 | 29 | 4.32 | 1.45 | 0.1789 | 29 | | | | |
| | **6M** | 13.70 | 4.81 | 0.0001 | 33 | 14.34 | 5.95 | <0.0001 | 33 | | | | |
| | **10M** | 20.96 | 7.17 | <0.0001 | 33 | 21.77 | 8.78 | <0.0001 | 33 | | | | |
| **WT** | **3M vs 6M** | 5.75 | 3.82 | 0.0022 | 16 | 5.55 | 3.97 | 0.0016 | 16 | | | | |
| | **3M vs 10M** | 0.50 | 0.36 | 0.7242 | 16 | 0.19 | 0.15 | 0.8810 | 16 | | | | |
| | **6M vs 10M** | -5.25 | 7.13 | <0.0001 | 17 | -5.36 | 7.58 | <0.0001 | 17 | | | | |
| **HET** | **3M vs 6M** | 16.70 | 12.05 | <0.0001 | 16 | 15.57 | 12.43 | <0.0001 | 16 | | | | |
| | **3M vs 10M** | 18.70 | 20.37 | <0.0001 | 16 | 17.64 | 20.11 | <0.0001 | 16 | | | | |
| | **6M vs 10M** | 2.01 | 1.97 | 0.0844 | 17 | 2.07 | 2.03 | 0.0749 | 17 | | | | |
