## Supplementary table 2 for "Mapping brain volume changes in the zQ175DN mouse model of Huntington’s disease: a longitudinal MRI study"

**Supplementary table 2:** Outcome of linear mixed model to test the main effects of age and genotype and (age x genotype) interaction on the absolute volumes of the cortex, striatum, corpus callosum and the ventricles ($p < 0.05$). Post hoc testing (FDR corrected, $p < 0.05$) was performed in case of significant (age x genotype) interaction. Red font indicates a significant effect.

| | Cortex | | | Striatum | | | Corpus callosum | | | Ventricles | | |
|---|---|---|---|---|---|---|---|---|---|---|---|---|
| **LMM effects** | **DF** | **F-value** | **P-value** | **DF** | **F-value** | **P-value** | **DF** | **F-value** | **P-value** | **DF** | **F-value** | **P-value** |
| **Genotype** | 1.56,51.39 | 113.3 | <0.0001 | 1.98,65.19 | 34.39 | <0.0001 | 1.96,64.58 | 0.1007 | 0.0002 | 1.70,56.16 | 91.7 | 0.3856 |
| **Age** | 1,34 | 50.23 | <0.0001 | 1,34 | 89.05 | <0.0001 | 1,34 | 17.63 | 0.9006 | 1,34 | 0.7725 | <0.0001 |
| **Genotype*age** | 2,66 | 24.07 | <0.0001 | 2,66 | 8.858 | 0.0004 | 2,66 | 3.271 | 0.0442 | 2,66 | 0.8815 | 0.4190 |

| | | Cortex | | | | Striatum | | | | Corpus callosum | | | | Ventricles | | | |
|---|---|---|---|---|---|---|---|---|---|---|---|---|---|---|---|---|---|
| **post-hoc effects** | | **mean dif** | **t-value** | **P-value** | **DF** | **mean dif** | **t-value** | **P-value** | **DF** | **mean dif** | **t-value** | **P-value** | **DF** | **mean dif** | **t-value** | **P-value** | **DF** |
| **WT -HET** | **3M** | 2.077 | 1.804 | 0.0810 | 30.99 | 0.9726 | 4.066 | 0.0005 | 29.88 | 0.3738 | 1.905 | 0.1707 | 31.53 | | | | |
| | **6M** | 4.824 | 6.534 | <0.0001 | 33.79 | 1.476 | 7.646 | <0.0001 | 24.52 | 0.5485 | 4.165 | 0.0010 | 32.71 | | | | |
| | **10M** | 8.708 | 10.13 | <0.0001 | 32.84 | 1.956 | 10.61 | <0.0001 | 25.56 | 0.8436 | 4.292 | 0.0010 | 33.99 | | | | |
| **WT** | **3M vs 6M** | 2.755 | 3.781 | 0.0021 | 16 | 0.3538 | 2.393 | 0.0330 | 16 | -0.1138 | 0.9523 | 0.4333 | 16 | | | | |
| | **3M vs 10M** | 3.981 | 6.2 | <0.0001 | 16 | 0.4519 | 3.114 | 0.0086 | 16 | -0.2217 | 1.703 | 0.1942 | 16 | | | | |
| | **6M vs 10M** | 1.225 | 2.565 | 0.0226 | 17 | 0.0981 | 0.496 | 0.6263 | 17 | -0.1078 | 0.8915 | 0.4333 | 17 | | | | |
| **HET** | **3M vs 6M** | 5.502 | 6.42 | <0.0001 | 16 | 0.8573 | 4.67 | 0.0005 | 16 | 0.06089 | 0.4934 | 0.6285 | 16 | | | | |
| | **3M vs 10M** | 10.61 | 12.19 | <0.0001 | 16 | 1.435 | 7.296 | <0.0001 | 16 | 0.2482 | 1.898 | 0.1707 | 16 | | | | |
| | **6M vs 10M** | 5.109 | 11.46 | <0.0001 | 17 | 0.5781 | 6.824 | <0.0001 | 17 | 0.1873 | 1.279 | 0.3271 | 17 | | | | |
