## Supplementary table 3 for "Mapping brain volume changes in the zQ175DN mouse model of Huntington’s disease: a longitudinal MRI study"

**Supplementary table 3:** Results of the linear mixed model to test the effects of age and genotype and (age x genotype) interaction on the log(Jacobian determinant) of the cortex, striatum, corpus callosum and the ventricles ROIs ($p < 0.05$). Red font indicates a significant effect.

| | Cortex | | | Cerebellum | | | Hippocampus | | |
|---|---|---|---|---|---|---|---|---|---|
| **LMM effects** | **DF** | **F-value** | **P-value** | **DF** | **F-value** | **P-value** | **DF** | **F-value** | **P-value** |
| **Genotype** | 1.75,57.71 | 534.3 | 0.0082 | 1.87,61.65 | 0.2506 | <0.0001 | 1.54,50.69 | 34.28 | 0.1719 |
| **Age** | 1,34 | 13.68 | 0.2191 | 1,34 | 133.5 | <0.0001 | 1,34 | 9.16 | 0.8079 |
| **Genotype*age** | 2,66 | 6.418 | <0.0001 | 2,66 | 89.1 | <0.0001 | 2,66 | 27.58 | <0.0001 |

| | Striatum | | | Globus pallidus | | | Substantia nigra | | |
|---|---|---|---|---|---|---|---|---|---|
| **LMM effects** | **DF** | **F-value** | **P-value** | **DF** | **F-value** | **P-value** | **DF** | **F-value** | **P-value** |
| **Genotype** | 1.49,49.07 | 99.95 | <0.0001 | 1.97,65.07 | 160.8 | <0.0001 | 1.64,54.15 | 172.2 | 0.0427 |
| **Age** | 1,34 | 185.2 | <0.0001 | 1,34 | 35.45 | <0.0001 | 1,34 | 4.435 | <0.0001 |
| **Genotype*age** | 2,66 | 3.09 | 0.0521 | 2,66 | 37.84 | <0.0001 | 2,66 | 10.99 | <0.0001 |
