## Supplementary table 4 for "Mapping brain volume changes in the zQ175DN mouse model of Huntington’s disease: a longitudinal MRI study"

**Supplementary table 4:** Post hoc testing (FDR corrected, $p < 0.05$) was performed in case of significant (age x genotype) interaction on the log(Jacobian determinant) of the cortex, cerebellum, hippocampus, striatum, globus pallidus, and substantia nigra. Red font indicates a significant effect.

| | | Cortex | | | | Cerebellum | | | | Hippocampus | | | |
|---|---|---|---|---|---|---|---|---|---|---|---|---|---|
| **post hoc effects** | | **mean dif** | **t-value** | **P-value** | **DF** | **mean dif** | **t-value** | **P-value** | **DF** | **mean dif** | **t-value** | **P-value** | **DF** |
| **WT -HET** | **3M** | 0.007 | 1.26 | 0.2191 | 27.33 | -0.034 | 5.62 | <0.0001 | 28.11 | 0.001 | 0.25 | 0.8079 | 26.79 |
| | **6M** | 0.012 | 2.83 | 0.0092 | 30.95 | -0.062 | 9.78 | <0.0001 | 25.53 | 0.006 | 1.40 | 0.2210 | 33.85 |
| | **10M** | 0.022 | 5.47 | <0.0001 | 33.97 | -0.091 | 16.66 | <0.0001 | 31.85 | 0.005 | 6.15 | <0.0001 | 33.81 |
| **WT** | **3M vs 6M** | 0.042 | 19.04 | <0.0001 | 16 | 0.013 | 3.22 | 0.0054 | 16 | -0.019 | 5.61 | 0.0001 | 16 |
| | **3M vs 10M** | 0.063 | 19.31 | <0.0001 | 16 | 0.026 | 6.72 | <0.0001 | 16 | -0.021 | 5.60 | 0.0001 | 16 |
| | **6M vs 10M** | 0.022 | 7.79 | <0.0001 | 17 | 0.013 | 5.76 | <0.0001 | 17 | -0.002 | 0.78 | 0.5012 | 17 |
| **HET** | **3M vs 6M** | 0.047 | 14.30 | <0.0001 | 16 | -0.015 | 6.47 | <0.0001 | 16 | -0.015 | 5.20 | 0.0002 | 16 |
| | **3M vs 10M** | 0.008 | 19.86 | <0.0001 | 16 | -0.031 | 13.57 | <0.0001 | 16 | 0.007 | 2.35 | 0.0476 | 16 |
| | **6M vs 10M** | 0.032 | 11.43 | <0.0001 | 17 | -0.015 | 5.31 | 0.0001 | 17 | 0.022 | 11.76 | <0.0001 | 17 |

| | | Striatum | | | | Globus pallidus | | | | Substantia nigra | | | |
|---|---|---|---|---|---|---|---|---|---|---|---|---|---|
| **post hoc effects** | | **mean dif** | **t-value** | **P-value** | **DF** | **mean dif** | **t-value** | **P-value** | **DF** | **mean dif** | **t-value** | **P-value** | **DF** |
| **WT -HET** | **3M** | | | | | -0.003 | 0.38 | 0.7085 | 28.3 | 0.005 | 0.21 | 0.8342 | 24.34 |
| | **6M** | | | | | -0.042 | 4.60 | 0.0001 | 33.83 | -0.007 | 4.16 | 0.0004 | 30.52 |
| | **10M** | | | | | -0.074 | 12.29 | <0.0001 | 29.63 | -0.040 | 2.19 | 0.0460 | 32.41 |
| **WT** | **3M vs 6M** | | | | | -0.024 | 5.04 | 0.0002 | 16 | -0.084 | 7.97 | <0.0001 | 16 |
| | **3M vs 10M** | | | | | -0.039 | 6.61 | <0.0001 | 16 | -0.122 | 9.50 | <0.0001 | 16 |
| | **6M vs 10M** | | | | | -0.015 | 3.31 | 0.0046 | 17 | -0.038 | 6.95 | <0.0001 | 17 |
| **HET** | **3M vs 6M** | | | | | -0.063 | 9.14 | <0.0001 | 16 | -0.160 | 16.05 | <0.0001 | 16 |
| | **3M vs 10M** | | | | | -0.109 | 18.44 | <0.0001 | 16 | -0.166 | 10.53 | <0.0001 | 16 |
| | **6M vs 10M** | | | | | -0.047 | 7.47 | <0.0001 | 17 | -0.007 | 0.52 | 0.6892 | 17 |
