## Supplementary figure 1 for "Mapping brain volume changes in the zQ175DN mouse model of Huntington’s disease: a longitudinal MRI study"

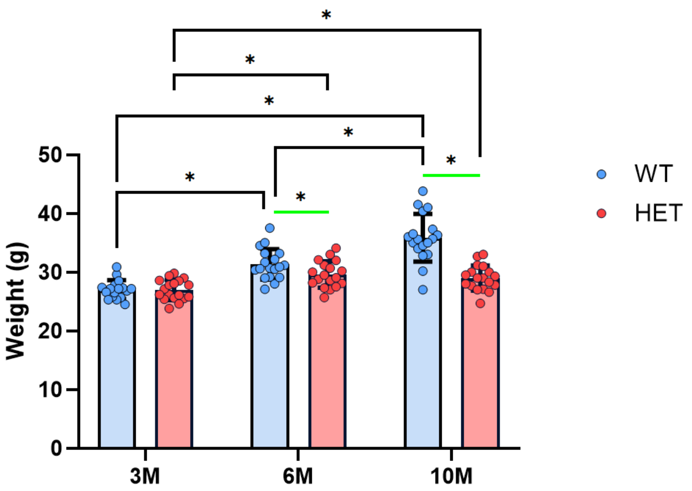


**Supplementary figure 1:** Comparison of body weight between zQ175DN HET and WT mice at 3M, 6M and 10M. Bar graph shows body weights for zQ175DN WT mice (blue) and HET mice (red). Black brackets indicate significant post hoc age by genotype effects, and green lines indicate significant genotype by age effects after FDR multiple comparisons correction. Error bars indicate SD. * $p < 0.05$.
