## Supplementary figure 2 for "Mapping brain volume changes in the zQ175DN mouse model of Huntington’s disease: a longitudinal MRI study"

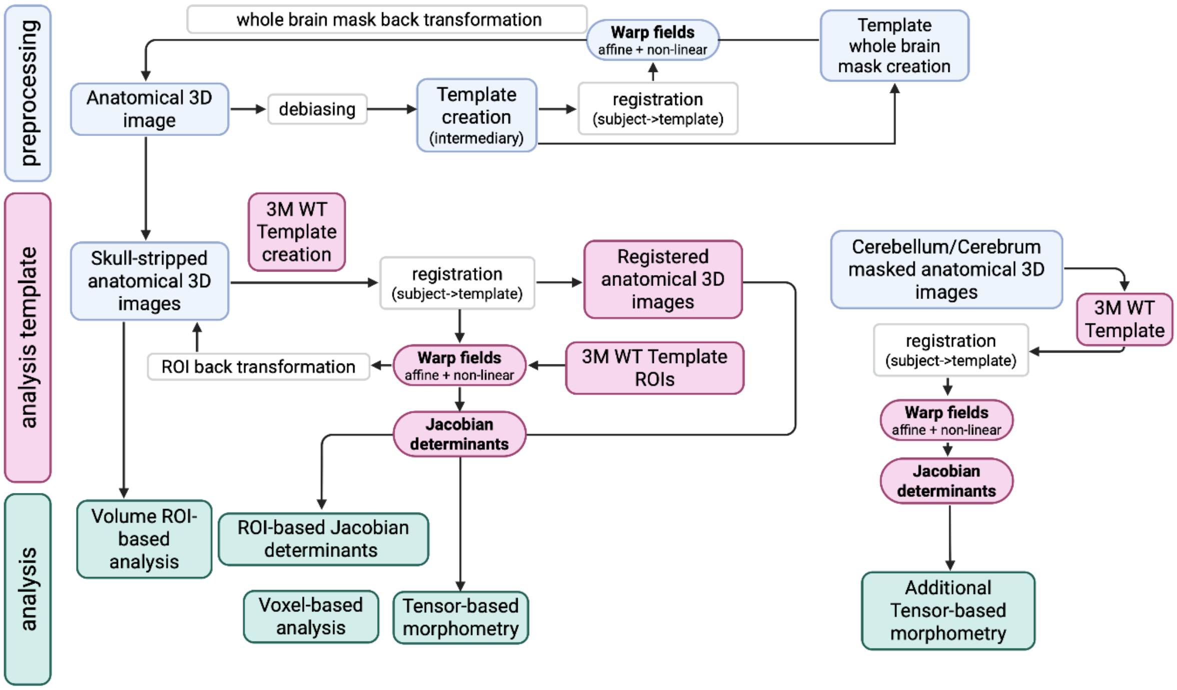


**Supplementary figure 2:** Flowchart summarizing the experimental pipeline for volumetric analysis and TBM analysis as described in the material and methods. Light blue boxes indicate preprocessing steps (debiasing, creating an intermediary template and skull stripping) and the output hereof. Pink boxes indicate processing steps required for creating the final analysis template (3M WT template) and Jacobian determinant warp fields. Green boxes indicate the analyses that were performed. These processing and analysis steps were repeated for the TBM analysis of cerebellum and cerebrum masked 3D images (right part flowchart).
