## Supplementary figure 3 for "Mapping brain volume changes in the zQ175DN mouse model of Huntington’s disease: a longitudinal MRI study"

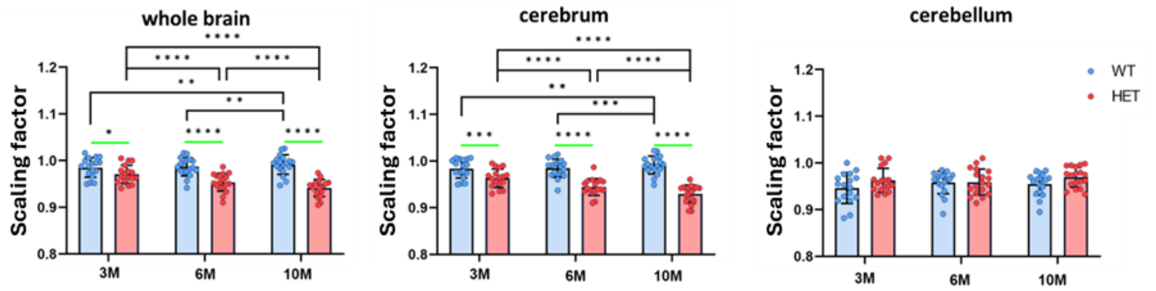


**Supplementary figure 3:** Scaling factors of whole, cerebrum and the cerebellum reflect absolute volume differences in HET mice. Bar graph of the affine scaling factors of whole brain, cerebrum and cerebellum used for TBM analysis. Bar graphs compare WT (blue) and HET (red) using a LMM with FDR correction. Green lines indicate post hoc genotype effects, black brackets indicate post hoc age effects. Horizontal dashed line indicates the main age effect. * $p < 0.05$, ** $p < 0.01$, **** $p < 0.0001$. Error bars represent *SD*.
