## Supplementary figure 4 for "Mapping brain volume changes in the zQ175DN mouse model of Huntington’s disease: a longitudinal MRI study"

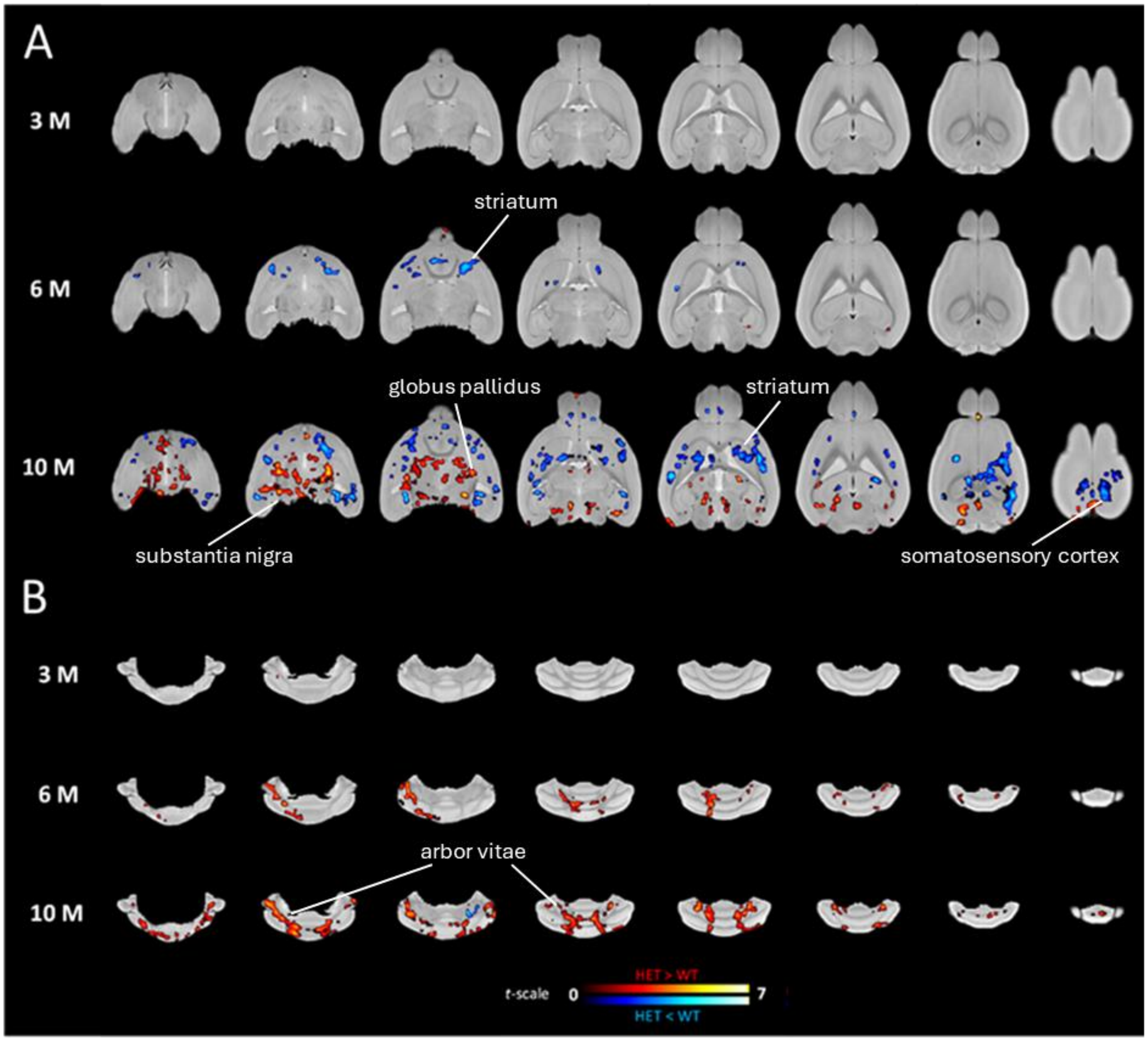


**Supplementary figure 4:** Tensor based morphometry showing age specific genotype differences using a cerebrum template-based TBM (A) or a cerebellum template-based TBM (B) of 3M wildtype subjects. **(A)** Statistical parametric maps showing voxels with a significant effect ($P_{TFCE\text{-}FWE} < 0.05$) in two two-sample t-tests comparing Jacobian determinants in the cerebrum between genotypes. Voxels with $P_{TFCE\text{-}FWE} < 0.05$ by age are overlaid on the 3M WT cerebrum template for 3M (top row), 6M (middle row) and 10M (bottom row). **(B)** Statistical parametric maps showing voxels with a significant effect ($P_{TFCE\text{-}FWE} < 0.05$) in two two-sample t-tests comparing Jacobian determinant in the cerebrum between genotypes. Voxels with $P_{TFCE\text{-}FWE} < 0.05$ by age are overlaid on the 3M WT cerebellum template for 3M (top row), 6M (middle row) and 10M (bottom row). Color bars indicate results of *t*-statistics with blue/red colors indicating significant voxels that have a lower relative log(Jacobian determinant)/higher relative log(Jacoban determinant) in HET compared to WT mice respectively.
